# Evolution of the α_2_-adrenoreceptors in vertebrates: ADRA2D is absent in mammals and crocodiles

**DOI:** 10.1101/106526

**Authors:** Héctor A. Céspedes, Kattina Zavala, Juan C. Opazo

**Author notes:** **Corresponding author:** Juan C. Opazo Instituto de Ciencias Ambientales y Evolutivas, Facultad de Ciencias, Universidad Austral de Chile, Valdivia, Chile.

## Abstract

Evolutionary studies of genes that have been functionally characterized and whose variation has been associated with pathological conditions represent an opportunity to understand the genetic basis of pathologies. α_2_-adrenoreceptors (ADRA2) are a class of G protein-coupled receptors that regulate several physiological processes including blood pressure, platelet aggregation, insulin secretion, lipolysis, and neurotransmitter release. This gene family has been extensively studied from a molecular/physiological perspective, yet much less is known about its evolutionary history. Accordingly, the goal of this study was to investigate the evolutionary history of α_2_-adrenoreceptors (ADRA2) in vertebrates. Our results show that in addition to the three well-recognized α_2_-adrenoreceptor genes (ADRA2A, ADRA2B and ADRA2C), we recovered a clade that corresponds to the fourth member of the α_2_-adrenoreceptor gene family (ADRA2D). We also recovered a clade that possesses two ADRA2 sequences found in two lamprey species. Furthermore, our results show that mammals and crocodiles are characterized by possessing three α_2_-adrenoreceptor genes, whereas all other vertebrate groups possess the full repertoire of α_2_-adrenoreceptor genes. Among vertebrates ADRA2D seems to be a dispensable gene, as it was lost two independent times during the evolutionary history of the group. Additionally, we found that most examined species possess the most common alleles described for humans; however, there are cases in which non-human mammals possess the alternative variant.

## 1. Introduction

Evolutionary studies of gene families that have been functionally characterized and whose variation has been linked to pathological conditions in humans represent an opportunity to understand the genetic basis of pathologies. Comparative studies have revealed that non-model species harbor genetic variation that could be relevant to understanding the physiological function of molecules that play key roles in health and disease (Opazo et al. 2005; Albertson et al. 2009; Yu et al., 2011; Edrey et al., 2012; Henning et al., 2014; Faulkes et al., 2015; Wichmann et al. 2016; Zavala et al., 2017). Studies carried out in non-model species that are resistant to diseases that have caused a high number of human deaths have been particularly important (e.g. Alzheimer, cancer)(Gorbunova et al., 2012; Castro-Fuentes and Socas-Pérez, 2013; Manov et al., 2013; Henning et al., 2014; Braidy et al., 2015; Faulkes et al., 2015; Inestrosa et al., 2015).

Adrenoceptors are a class of G protein-coupled cell membrane receptors that are mediators in the sympathetic nervous system. They regulate physiological functions to maintain homeostasis by mediating the action of catecholamines such as epinephrine and norepinephrine. There are two groups of adrenergic receptors, α and β, and there are several subtypes in each group. Within the α-type, the α_2_-adrenoreceptors (ADRA2) are composed of three subtypes: ADRA2A, ADRA2B and ADRA2C (Bylund et al., 1988). They are mainly expressed in the nervous system and platelets and regulate several key physiological processes including blood pressure, platelet aggregation, insulin secretion, lipolysis, and neurotransmitter release (Knaus et al. 2007). They also play significant roles during development as the deletion of the three α_2_-adrenoreceptor genes results in embryonic lethality (Philipp et al., 2002). At the individual level, the α_2A_-adrenoreceptors (ADRA2A) mediate inhibition of insulin release (Gribble, 2010; Rosengren et al., 2010), hypotension, bradycardia, baroreceptor reflex sensitivity, sedation, and hypnosis (MacMillan et al., 1996; Lakhlani et al., 1997, Niederhoffer et al., 2004). The α_2B_-adrenoreceptors (ADRA2B) play important roles in the regulation of vascular tone (Link et al.,1996) as well as in the development of the placenta (Muthig et al., 2007) and lungs (Haubold et al., 2010). The α_2C_-adrenoreceptors (ADRA2C) control the secretion of epinephrine via an autocrine feedback loop in the adrenal medulla (Brede et al., 2002, 2003; Gilsbach et al., 2007). In the past, a fourth type of ADRA2 receptor was described (ADRA2D) based on pharmacological assays (Bylund, 2005). This fourth type of receptor was initially identified in rat, mouse, and cow and was mainly characterized as having a low affinity for yohimbine and rauwolscine. However, subsequent studies have revealed that this receptor in these species would not be a fourth type of α_2_-adrenoreceptor; instead it would be an ADRA2A type of receptor with divergent pharmacological properties (Lanier et al., 1991; Link et al., 1992). Thus, α_2_-adrenoreceptor gene family is a group of cell membrane receptors that plays significant physiological roles from embryogenesis to adulthood that would be greatly benefited of having evolutionary information.

Accordingly, the main goal of this study was to unravel the evolutionary history of α_2_-adrenoreceptors (ADRA2) in vertebrates. To do so we annotated α_2_-adrenoreceptor (ADRA2) genes in species representative of all main groups of vertebrates. Using phylogenetic and syntenic approaches, we defined the composition of the gene family, orthologous relationships within each family member, and patterns of differential retention. We also studied genetic variability in non-model species related to humans; specifically, we analyzed variability at sites known to have consequences in human health. Our results show that in addition to the three well-recognized α_2_-adrenoreceptor genes (ADRA2A, ADRA2B and ADRA2C), we recovered a clade that corresponds to the fourth member of the α_2_-adrenoreceptor gene family (ADRA2D). We also recovered a clade that possesses two α_2_-adrenoreceptor sequences found in two species of lampreys. Based on the phyletic distribution of the genes, we show that mammals and crocodiles are characterized by possessing three α_2_-adrenoreceptor genes, whereas all other vertebrate groups possess the full repertoire of four α_2_-adrenoreceptor genes. The most dispensable α_2_-adrenoreceptor gene seems to be the ADRA2D gene as it was lost two independent times during the evolutionary history of vertebrates. We found that most examined species possess the most common alleles described for humans. There is a group of sites for which some species possess the most common allele, whereas others possess the alternative allele. There is one position for which the rare allele is present in all non-human species, and there are some cases in which novel alleles are present.

## 2. Materials and methods

### 2.1 DNA data and phylogenetic analyses

We used bioinformatic procedures to annotate α_2_-adrenoreceptor genes in species of all major groups of vertebrates. Our sampling included species from mammals, birds, reptiles, amphibians, coelacanths, teleost fish, holostean fish, cartilaginous fish and cyclostomes (Supplementary Table S1). We also included sequences of the dopamine receptors D (DRD) 1, 2, 3, 4 and 5, and β-adrenoreceptors (ADRB) 1, 2 and 3 from humans (Supplementary Table S1). Amino acid sequences were aligned using the FFT-NS-i strategy from MAFFT v.6 (Katoh and Standley, 2013). We used the proposed model tool of IQ-Tree (Trifinopoulos et al., 2016) to select the best-fitting model of amino acid substitution (JTT + R6). Phylogenetic relationships were estimated according to maximum likelihood approach. We performed a maximum likelihood analysis to obtain the best tree using the program IQ-Tree (Trifinopoulos et al., 2016) and assessed support for the nodes with 1,000 bootstrap pseudoreplicates using the ultrafast routine. Human ADRA1A, B, and D sequences were used as outgroups.

### 2.2 Assessments of Conserved Synteny

We examined genes found upstream and downstream of the α_2_-adrenoreceptor genes of species representative of all main groups of vertebrates. We used the estimates of orthology and paralogy derived from the EnsemblCompara database (Herrero et al., 2016); these estimates are obtained from an automated pipeline that considers both synteny and phylogeny to generate orthology mappings. These predictions were visualized using the program Genomicus v86.01 (Louis et al., 2015). Our analyses were performed in humans (*Homo sapiens*), opossum (*Monodelphis domestica*), chinese turtle (*Pelodiscus sinensis*), chicken (*Gallus gallus*), anole lizard (*Anolis carolinensis*), clawed frog (*Xenopus tropicalis*), coelacanth (*Latimeria chalumnae*), spotted gar (*Lepisosteus oculatus*) and elephant shark (*Callorhinchus milii*). In the case of the elephant shark (http://esharkgenome.imcb.a-star.edu.sg/), the genomic pieces containing *ADRA2* genes were annotated, and predicted genes were then compared with the non-redundant protein database using Basic Local Alignment Search Tool (BLAST) (Altschul et al., 1990).

## 3. Results and discussion

### 3.1. Gene trees, synteny analyses, and orthology

We constructed a phylogenetic tree in which we included ADRA2 sequences of representative species of all major groups of vertebrates. Our phylogenetic analyses recovered the monophyly of the ADRA2 sequences with strong support (Fig. 1). Within the ADRA2 clade, we recovered five monophyletic groups that correspond to the three already well characterized α_2_-adrenoreceptor genes (ADRA2A, ADRA2B and ADRA2C), a clade that correspond to the fourth member of the α_2_-adrenoreceptor gene family (ADRA2D), plus a clade that contains the ADRA2 sequences of two lamprey species (Fig. 1). It is important to note that the ADRA2D clade we identified in our phylogenetic analyses does not correspond to the clade that includes the α_2_-adrenoreceptor gene of rat, mouse and cow that in the past was named ADRA2D (Bohmann et al. 1994; Kobayashi et al. 2004; Bylund, 2005). The use of the name ADRA2D to refer to the clade including the α_2_-adrenoreceptor gene of rat, mouse, and cow was assigned based on pharmacological assays; specifically, lower affinity of yohimbine and rauwolscine for the receptor. Homology, however, is an evolutionary concept that is related but not based on functionality (Gabaldón, 2008; Gabaldón & Koonin, 2013). Subsequent studies have revealed that what was called α_2D_-adrenoreceptor (ADRA2D) in rat and mouse in actuality is not a fourth type of α_2_-adrenoreceptor; instead it is an ortholog to the α_2A_-adrenoreceptor (ADRA2A) of other mammals (Lanier et al., 1991; Link et al., 1992).

**Figure 1.**
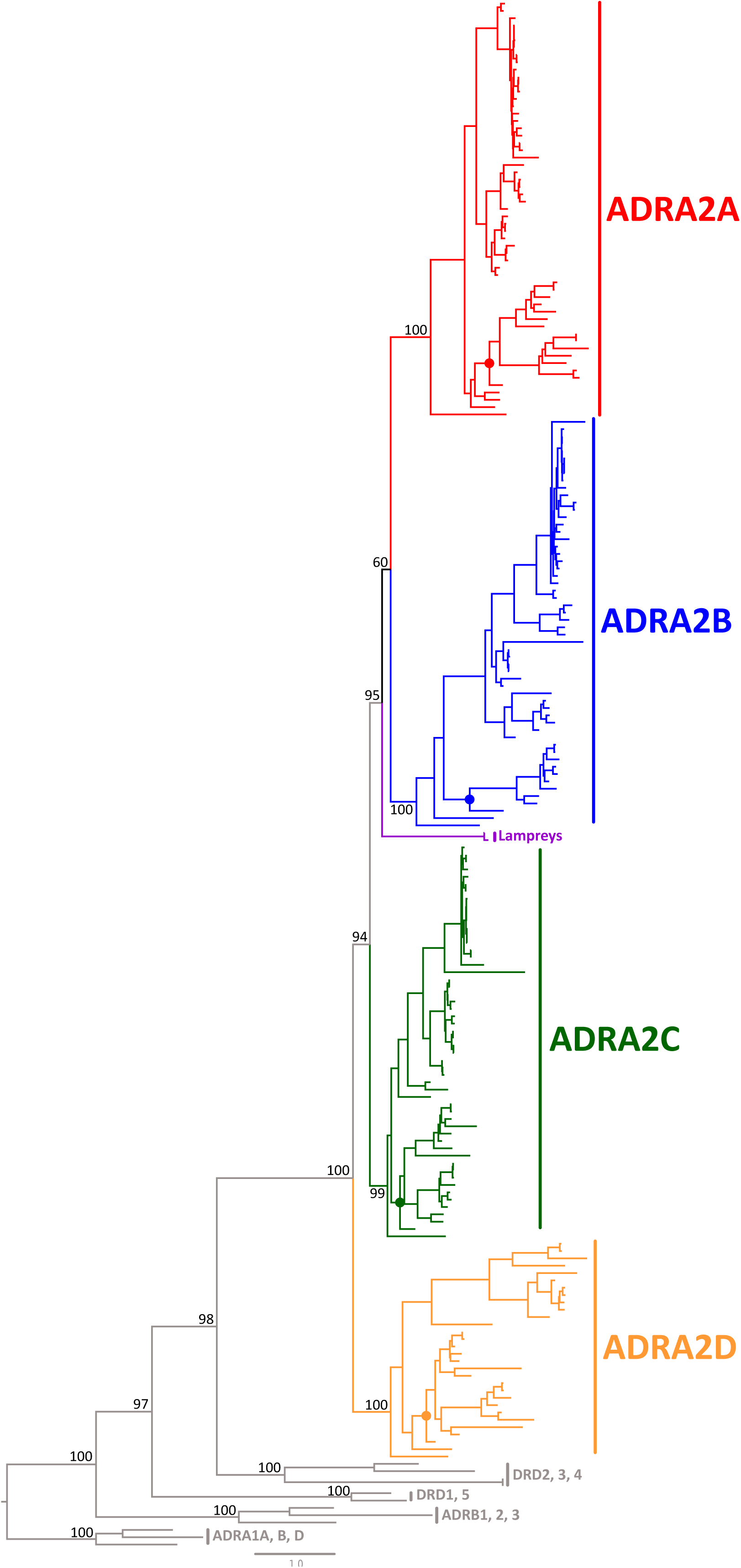
Maximum likelihood tree depicting evolutionary relationships among α_2_-adrenoreceptors. Numbers on the nodes correspond to maximum likelihood ultrafast bootstrap support values. Human ADRA1A, B, and D sequences were used as outgroups. Color dots denote the fish subtrees that are expanded in figure 4.

Within the ADRA2 clade, phylogenetic relationships among different members of the gene family are also well supported (Fig. 1). We recovered a monophyletic group containing the ADRA2A and lamprey clades, which in turn was recovered as sister to the ADRA2B sequences (Fig. 1). The monophyletic group including ADRA2C sequences was recovered as sister to the aforementioned clade (Fig.1), whereas the ADRA2D clade was recovered as sister to all other ADRA2 groups (Fig. 1). In agreement with previous results, the clade that includes dopamine receptors (DRD) 2, 3 and 4 was recovered as sister to the ADRA2 clade (Spielman et al., 2015; Zavala et al., 2017). Although we expected dopamine receptors (DRD) 1 and 5 to be sister to the β-adrenoreceptors (ADRB) (Spielman et al., 2015; Zavala et al., 2017), this was not the case. This could mainly be due to the fact that the sampling effort performed in this study was not aimed at resolving the evolutionary relationship among these genes. The synteny analyses gave further support for the identity of the four α_2_-adrenoreceptor clades identified for all of the main groups of gnathostomes (Fig. 2). Although we found variation in the pattern of conservation of genes found up-and downstream of the ADRA2 genes, in all cases it is possible to realize that each ADRA2 gene is located in a chromosomal location that is more or less conserved in all examined species (Fig. 2). For example, in the case of the ADRA2A gene, there are four upstream genes (SHOC2, BBIP1, PDCD4 and RBM20) and five downstream genes (GPAM, TECTB, ACSL5, ZDHHC6 and VTI1A) that are well conserved in most examined species (Fig. 2); this suggests that this gene arrangement was present in the common ancestor of the group between 615 and 473 million years ago and was inherited by all descendant lineages. It is worth noting that the genomic region where the ADRA2D gene is located is also conserved in species that lost the gene (e.g. humans; Fig. 2).

**Figure 2.**
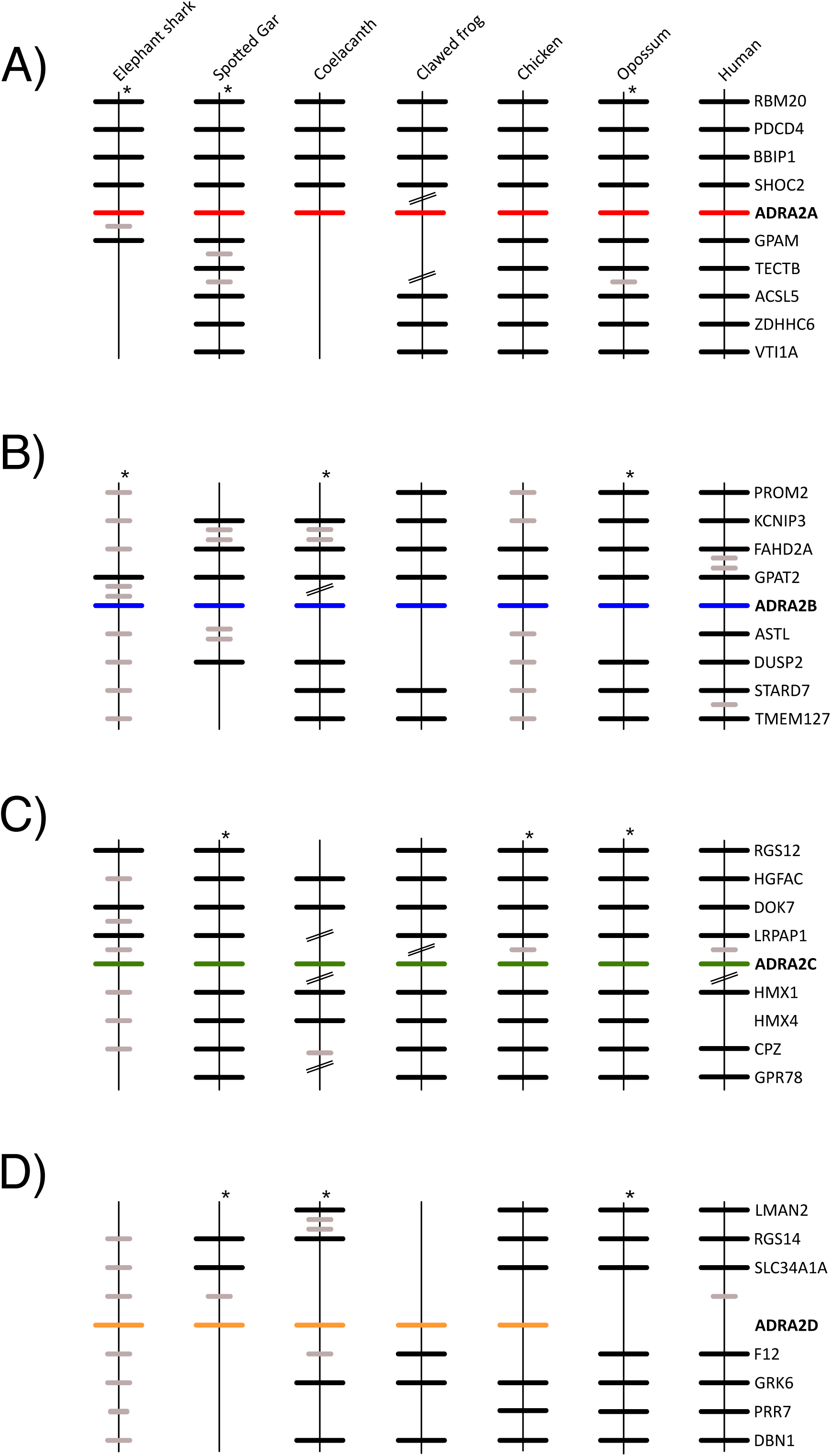
Conserved synteny in the chromosomal regions that harbor α_2_-adrenoreceptor genes. A) Chromosomal region that harbors ADRA2A genes; B) region that harbors the ADRA2B genes; C) Chromosomal region that harbors the ADR2C genes; D) Chromosomal region that harbors the ADR2D genes. Asterisks denote that the orientation of the genomic piece is from 3´to 5´.

From the phylogenetic analyses it was possible to define orthology among the gnathostome sequences, however, this was not possible for the lamprey sequences (Fig. 1). Genome-wide compositional analyses have shown that lamprey genomes possess compositional bias (Qiu et al., 2011; Kuraku, 2013; Mehta et al., 2013; Smith et al., 2013) that makes it difficult to recover the true evolutionary history of genes using phylogenetic inference. In support of this idea, other studies working with different gene families have also observed that lamprey sequences tend to group together instead of with their true ortholog (Qiu et al., 2011; Nah et al., 2014; Campanini et al., 2015; Opazo et al., 2015; Wichmann et al., 2016; Zavala et al., 2017). In the case of the sea lamprey, we also explored the genomic piece in which we found the ADRA2 gene to determine if syntenic genes could be found that would help us to define orthology. In principle, if we found syntenic genes in lampreys that have a human ortholog located in one of the chromosomes where the ADRA2 genes are located, we could suggest orthology. In our analyses, we found two genes (RNF122 and C20orf27) on the 3´ side of the sea lamprey ADRA2 gene; unfortunately, neither of these genes had an ortholog on a human chromosome where ADRA2 genes are located. The RNF122 gene is located on human chromosome 8, while C20orf27 is located on human chromosome 20.

### 3.2. Diversification and differential retention of ADRA2 genes during the evolutionary history of vertebrates

In this study we identified four types of α_2_-adrenoreceptors in species representative of all of the main groups of vertebrates (ADRA2A, ADRA2B, ADRA2C and ADRA2D; Fig. 1). The phyletic distribution of the α_2_-adrenoreceptors found here suggests that the last common ancestor of the group, which existed between 676 and 615 million years ago, possessed the full repertoire of α_2_-adrenoreceptors observed in extant species. However, given that the last common ancestor of vertebrates underwent two rounds of whole genome duplication (Ohno et al., 1968; Meyer and Schartl, 1999; McLysaght et al., 2002; Dehal and Boore, 2005; Hoegg and Meyer, 2005; Putnam et al., 2008), it is possible that at some point in time the vertebrate ancestor had just one ADRA2 gene, which after the two rounds of whole genome duplication gave rise to the actual repertoire of four genes. In support of this idea, the α_2_-adrenoreceptor gene family appears in the repository of genes that were retained after the whole genome duplications occurred in the vertebrate ancestor (Singh et al., 2015). This pattern of gene diversification has also been observed for other gene families (Blomme et al., 2006; Hoffmann and Opazo, 2011; Hoffmann et al., 2012; Storz et al., 2013; Wichmann et al., 2016; Zavala et al., 2017), highlighting the pivotal role of whole genome duplication in the origin of biological diversity.

After the repertoire of four α_2_-adrenoreceptor genes was established, vertebrate groups retained different complements of α_2_-adrenoreceptor genes (Fig. 3). According to our assessment, all but two groups retained the full complement of α_2_-adrenoreceptor genes (Fig. 3). Only mammals and crocodiles retained three family members (Fig. 3), and in both cases the lost gene was the α_2D_-adrenoreceptor gene (ADRA2D; Fig. 3) which is likely a dispensable gene as it has been lost two independent times during the evolutionary history of vertebrates (Fig. 3). Based on the phyletic distribution of the α_2_-adrenoreceptor genes, it is possible to suggest that for mammals and crocodiles just having three members of the gene family is enough to maintain all biological functions associated with this group of genes. Likewise it is possible that there is some degree of redundancy that works as a backup (i.e. functionally overlapping paralogues) in the event that one of the genes is lost or inactivated (Gitelman, 2007; Cañestro et al., 2009; Félix & Barkoulas, 2015; Albalat & Cañestro, 2016). Thus, the differential retention of α_2_-adrenoreceptor genes could be seen as a stochastic process, in which the observed differences in gene repertoires do not necessarily translate into functional consequences. However, it is also possible that the retention of multiple (redundant) gene copies could help to direct the trajectory of physiological evolution by providing an increased opportunity for the origin of biological novelties (Ohno et al., 1968; Ohno, 1970; Force et al., 1999; Hughes, 1994; Zhang, 2003).

**Figure 3.**
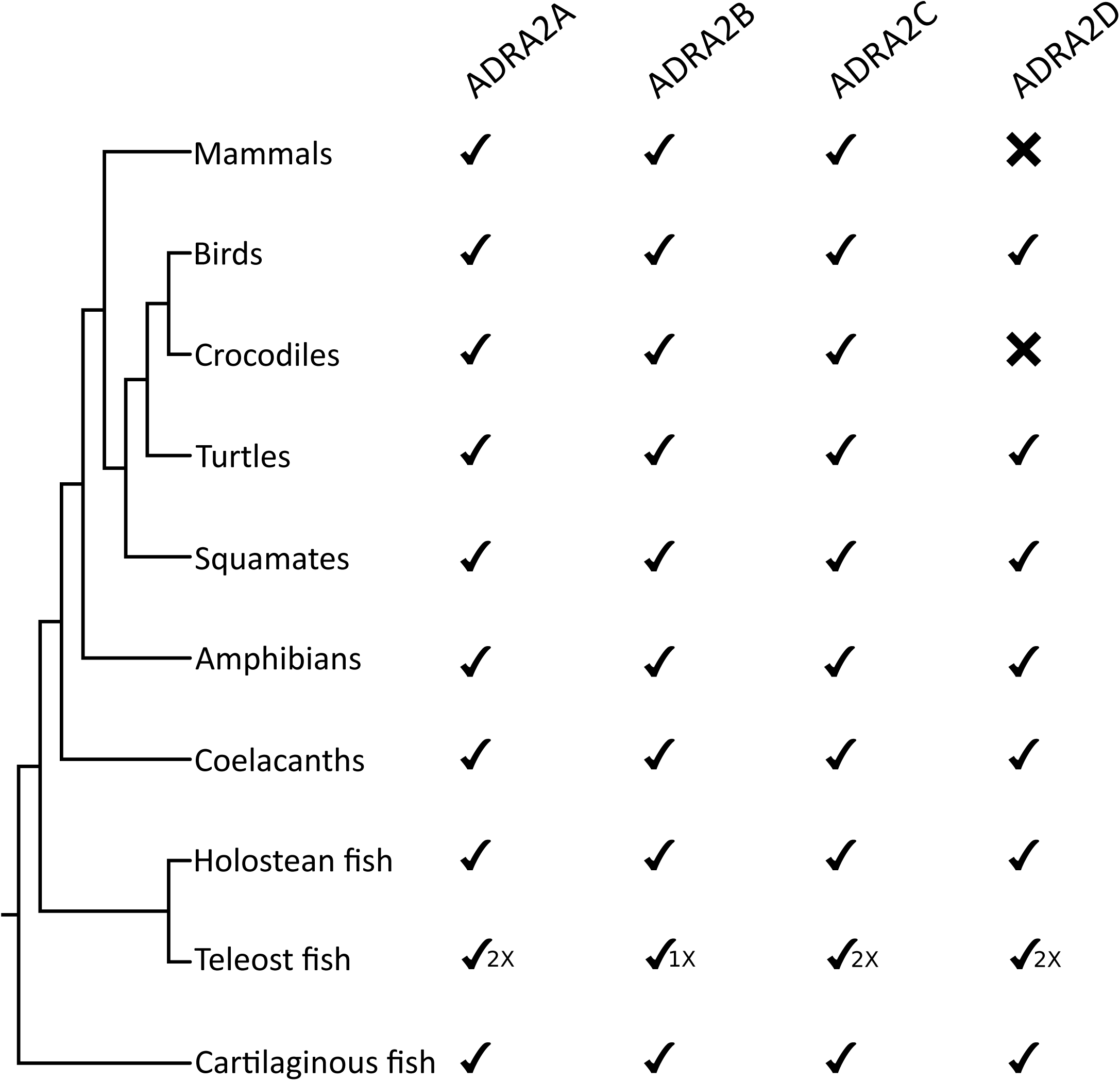
Phyletic distribution of α_2_-adrenoreceptor genes in gnathostomes.

The diversification of the α_2_-adrenoreceptor genes in teleost fish deserves a special mention given that the last common ancestor of this group underwent an extra round of whole genome duplication in comparison to other vertebrates (Meyer and Van de Peer, 2005; Kasahara, 2007; Sato and Nishida, 2010). Given this, one could expect to find duplicated copies of all α_2_-adrenoreceptor gene lineages in teleost fish compared to the spotted gar, a holostean fish species that did not experience the extra round of whole genome duplication. In this case, special attention should be paid to synteny, given that the asymmetric loss of duplicates could cause one to misinterpret paralogy as orthology i.e. hidden paralogy (Daubin et al., 2001; Martin & Burg, 2002; Kuraku, 2013; Kuraku et al., 2016). According to our phylogenetic tree, duplicated lineages were found in three (ADRA2A, ADRA2C and ADRA2D) out of four α_2_-adrenoreceptor genes (Fig. 4). In the case of ADRA2B, our phylogenetic analyses allowed us to identify one α_2_-adrenoreceptor gene lineage in nine species of teleost fish (Fig. 4). Synteny analyses provided extra support for the identity of this group of sequences as a gene lineage as we identified two genes upstream (GPAT2 and FAHD2A) and one downstream (DUSP2) that are conserved in most species. The presence of a single ADRA2B gene lineage in this group of teleost fish suggests that the loss of others occurred early in the evolutionary history of teleost fish.

**Figure 4.**
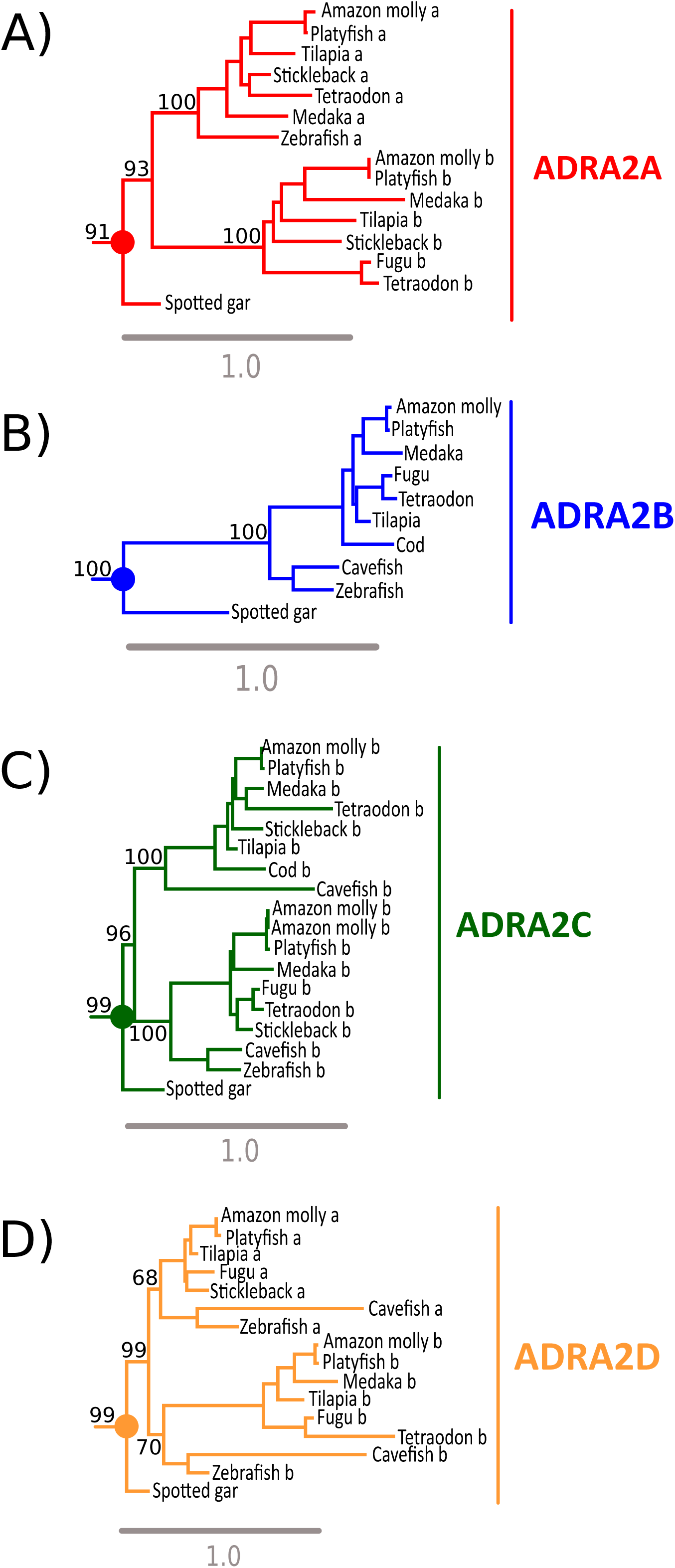
Maximum likelihood trees depicting evolutionary relationships among α_2_-adrenoreceptors in fish. A) phylogenetic relationships among ADRA2A sequences; B) phylogenetic relationships among ADR2B sequences; C) phylogenetic relationships among ADRA2C sequences; D) phylogenetic relationships among ADRA2D sequences. Numbers on the nodes correspond to maximum likelihood ultrafast bootstrap support values. These tree topologies do not represent novel phylogenetic analyses; they are the fish clades that were recovered from figure 1.

### 3.3 α_2_-adrenoreceptor polymorphisms in species related to humans

Although there are several polymorphisms described for the α_2_-adrenoreceptor genes, in this work we only studied the variants that have been functionally characterized (Ahles and Engelhardt, 2014). Screening species closely related to humans could potentially shed light on the functional role of α_2_-adrenoreceptor gene variation. Especially important are the cases in which all closely related species possess the (alternative) allele that is associated with given pathological conditions in humans. This situation could indicate that the specific position of the allele itself does not produce the human pathological condition, rather pathology depends on the identity at other amino acid positions within the molecule (Breen et al., 2012; Natarayan et al., 2013).

In our survey, we compared polymorphisms that have been described and functionally characterized in humans in a panel of species that includes anthropoid primates and rodents (Table 1). We found that most examined species possess the most common alleles described for humans (Table 1). There is also a group for which some species possess the most common allele, whereas others possess the alternative allele (Table 1). Additionally, we found one position for which the rare allele is present in all non-human species (Table 1), and there are some cases in which novel alleles are present (Table 1). The most interesting case is the polymorphism located approximately 70 kb (rs869244) downstream of the ADRA2A locus (Table 1). This polymorphism has been associated with platelet aggregation, a physiological response to vessel injury in response to three different agonists (ADP, epinephrine and collagen) (Johnson et al., 2010). Among humans and considering all human populations together, the alternative allele (adenine) is present at a frequency of 38%; the frequency of this allele varies from 32% in African populations to 44% in South Asian populations (Kohler et al., 2014). According to our results, all non-human species possess the alternative allele (Table 1). Although it is possible that in non-human mammals the frequency of the minor allele follows the human trend, and we only screened individuals belonging to the 38% of the population, the fact that all examined species possess the alternative allele would suggest that adenine at this position is common in non-human mammalian species. If that were the case, our data would suggest that the specific position of the allele by itself potentially does not produce the human pathological condition; rather pathology depends on the identity at other positions.

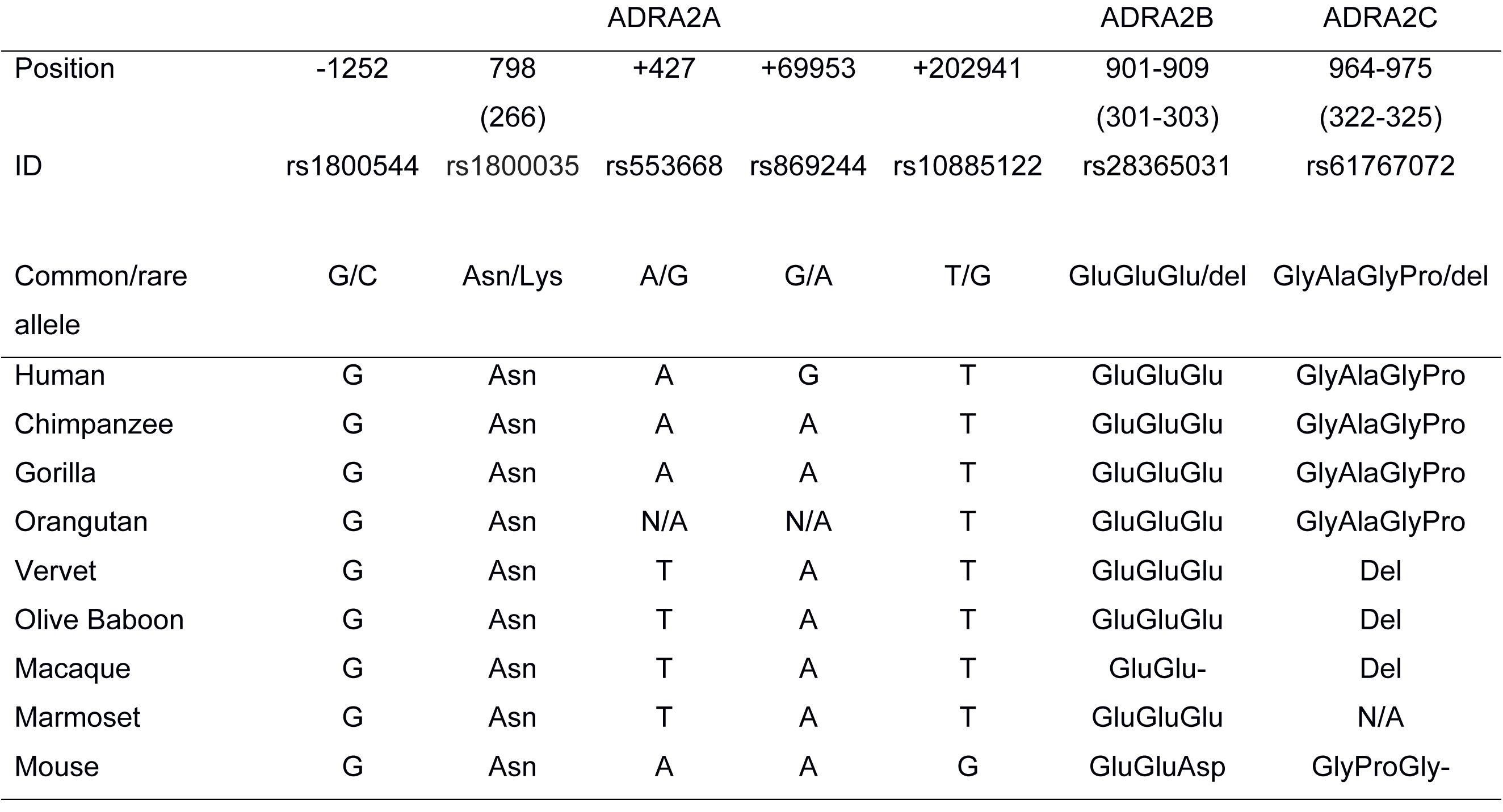

|  | ADRA2A |  |  |  |  | ADRA2B | ADRA2C |
| --- | --- | --- | --- | --- | --- | --- | --- |
| Position | -1252 | 798 | +427 | +69953 | +202941 | 901-909 | 964-975 |
|  |  | (266) |  |  |  | (301-303) | (322-325) |
| ID | rs1800544 | rs1800035 | rs553668 | rs869244 | rs10885122 | rs28365031 | rs61767072 |
| Common/rare allele | G/C | Asn/Lys | A/G | G/A | T/G | GluGluGlu/del | GlyAlaGlyPro/del |
| Human | G | Asn | A | G | T | GluGluGlu | GlyAlaGlyPro |
| Chimpanzee | G | Asn | A | A | T | GluGluGlu | GlyAlaGlyPro |
| Gorilla | G | Asn | A | A | T | GluGluGlu | GlyAlaGlyPro |
| Orangutan | G | Asn | N/A | N/A | T | GluGluGlu | GlyAlaGlyPro |
| Vervet | G | Asn | T | A | T | GluGluGlu | Del |
| Olive Baboon | G | Asn | T | A | T | GluGluGlu | Del |
| Macaque | G | Asn | T | A | T | GluGlu- | Del |
| Marmoset | G | Asn | T | A | T | GluGluGlu | N/A |
| Mouse | G | Asn | A | A | G | GluGluAsp | GlyProGly- |

The polymorphism described for the ADRA2C gene represents an inframe deletion of four amino acids (GlyAlaGlyPro), between positions 322 and 325, in the third intracellular loop. Some studies have associated this polymorphism, in combination with the identity of position 389 (Arg) of the ADRB1 gene, with the risk for cardiovascular events (e.g. stroke, cardiac death) and early onset of hypertension (Small et al., 2002). However, there are other studies in which this association is unclear (Nonen et al., 2005; Metra et al., 2006; Canham et al., 2007). In our study, we found that all examined Old World monkeys (vervet, olive baboon, and macaque) possess the alternative allele, i.e a deletion of four amino acids (Table 1), suggesting that this variant could be common in this group of primates. Furthermore, according to Zavala et al. (2017), in Old World monkeys arginine at position 389 of the ADRB1 gene is also common. Thus, Old World monkeys possess both variants that have been associated with the risk of cardiovascular disease and early onset hypertension (Small et al., 2002). However, similarly to what was mentioned for the ADRA2A polymorphism, it is possible that genetic variants by themselves do not trigger the condition to which they have been linked; their effects could depend on the identity at other amino acid positions.

## 4. Conclusions

In this study, we present an evolutionary analysis of the α_2_-adrenoreceptor gene family in representative species of vertebrates. Our phylogenetic and synteny analyses show that in addition to the three well-recognized α_2_-adrenoreceptor genes (ADRA2A, ADRA2B and ADRA2C), we also recovered a clade that corresponds to the fourth member of the α_2_-adrenoreceptor gene family (ADRA2D). Based on the phyletic distribution of the genes, we show that all but two vertebrate groups retained the full complement of α_2_-adrenoreceptor genes; mammals and crocodiles are characterized by possessing three α_2_-adrenoreceptor family members. Furthermore, the α_2D_-adrenoreceptor could be a dispensable gene as it was lost two independent times during the evolutionary history of vertebrates. The pattern of differential retention observed for this group of genes opens new avenues to study the biology of the different α_2_-adrenoreceptors. For example, mammals and crocodiles could be seen as natural knockout models for the α_2D_-adrenoreceptor gene (ADRA2D). In addition to the above, studies of species that possess the full complement of α_2_-adrenoreceptors (e.g. spotted gar) would provide significant evolutionary information regarding the division of labor in the α_2_-adrenoreceptor gene family.

## Acknowledgments

This work was funded by Fondo Nacional de Desarrollo Científico y Tecnológico (FONDECYT) grant 1160627 to JCO.

## Appendix A. Supplementary data

Supplementary data associated with this article can be found in the online version.

